# Identification and Description of Emotions by Current Large Language Models

**DOI:** 10.1101/2023.07.17.549421

**Authors:** Suketu C. Patel, Jin Fan

## Abstract

The assertion that artificial intelligence (AI) cannot grasp the complexities of human emotions has been a long-standing debate. However, recent advancements in large language models (LLMs) challenge this notion by demonstrating an increased capacity for understanding and generating human-like text. In this study, we evaluated the empathy levels and the identification and description of emotions by three current language models: Bard, GPT 3.5, and GPT 4. We used the Toronto Alexithymia Scale (TAS-20) and the 60-question Empathy Quotient (EQ-60) questions to prompt these models and score the responses. The models’ performance was contrasted with human benchmarks of neurotypical controls and clinical populations. We found that the less sophisticated models (Bard and GPT 3.5) performed inferiorly on TAS-20, aligning close to alexithymia, a condition with significant difficulties in recognizing, expressing, and describing one’s or others’ experienced emotions. However, GPT 4 achieved performance close to the human level. These results demonstrated that LLMs are comparable in their ability to identify and describe emotions and may be able to surpass humans in their capacity for emotional intelligence. Our novel insights provide alignment research benchmarks and a methodology for aligning AI with human values, leading toward an empathetic AI that mitigates risk.

## Introduction

Many researchers have challenged the possibility of artificial intelligence (AI) systems’ ability to exhibit empathy, emotional intelligence, and social and physical understanding [1,2,3]. Frameworks for developing emotional machines have proposed that embodiment is required [4,5,6]. It has been contended that AI will never truly comprehend emotions as they are subjective, private, and often fleeting, positing that the elusive nature of emotions is incompatible with the objectivity inherent in AI systems [7]. Similarly, others have echoed this sentiment, asserting that AI’s incapacity for subjective experiences limits its understanding and response to emotions, thereby restricting it from attaining true intelligence or developing genuine empathy [2]. A recent analysis has detailed the inherent limitations of empathy in AI, stating that these systems lack both cognitive and affective components necessary for empathetic experiences [. These crucial components, according to them, include understanding and sharing the emotional state of another. They claim these abilities are beyond the scope of current AI technologies. From a neurobiological perspective, the multifaceted nature of emotions and their connection to various brain systems present an insurmountable challenge for AI [9]. Another known criticism of AI argues that the unique combination of human intelligence and life experiences is an aspect that machines cannot replicate, hence making emotional understanding an unattainable goal for AI systems [10].

Others have predicted that “by 2029, computers will have emotional intelligence and be convincing as people” [11] citing the exponential growth trajectory of computing power and information technology. Traditional AI methods, primarily algorithm-based, have yet to advance our understanding of emotion significantly. However, there are promising unified theories of high-level intelligence that incorporate both emotional and embodied intelligence [12,5]. Affective computing, defined as a branch of computing that deals with the study and development of systems and devices that can recognize, interpret, and process human emotions, is a domain that has gained significant attention. Research is increasingly focusing on imbuing AI with artificial empathy [13], an aspect of affective computing that involves the ability of an AI system to understand and respond to human emotions. This strategy expands the boundaries of cognitive and emotional processing capabilities within AI systems [14,15]. While artificial emotional intelligence [16] refers to the capability of a machine to recognize and respond to emotions, similar to how humans do, artificial empathy [17] is specifically about enabling machines to understand and respond to human emotional states in a manner that exhibits empathy. Although both focus on machine understanding of human emotions, the key difference lies in their application; the former is about broad emotional responsiveness, while the latter concentrates on empathetic understanding and response. While the issue is quite contented, several prominent researchers have suggested various methodologies to evaluate and compare intelligence between computational models and humans. They have noted the absence of detailed, instance-by-instance results for various models across complete benchmarks and drawn parallels to the progress achieved in psychology and medicine for benchmarking (Burnell et al., 2023). The ultimate goal is to match or surpass these advancements in the context of intelligence models [18, 19].

If the AI system does not show a human level of emotional processing [20], it can have impairments with varying degrees of alexithymia traits. These traits present as challenges in identifying and expressing one’s emotions, recognizing emotions in others, and overall emotional intelligence [21], which refers to a deficiency in the experience and processing of emotions [22]. The meaning of alexithymia is “lack of words for emotion,” and also includes externally-oriented thinking. An externally-oriented thinking style alludes to an individual’s propensity towards concrete reasoning, being influenced by immediate stimuli, and focusing on the pragmatic facets of a given situation [23]. People with alexithymia may struggle to articulate their feelings and appear emotionally distant, and alexithymia has been widely researched in the field of psychology and psychiatry [22]. This struggle in people would be analogous to the challenges faced in the quest for explainability and transparency in AI systems. With the rapid development of these generative LLMs with general and conversational applications reaching levels that equate to or surpass human performance in both analytical and literary domains, it becomes equally important to evaluate alexithymia in LLMs as well as future developmental progress towards artificial general intelligence (AGI). Its clinical applications show that LLM cognitive language capacity is advanced enough to predict dementia from speech [24]. Historically, the diagnosis of alexithymia has been reserved for humans. However, the emergence of LLMs [25] such as ChatGPT has paved the way for new avenues of exploration. It has now become possible to study this trait within language models. These models have learning systems that trace their roots back to the architecture of human neural networks [26] Over the years, with the improvement in computational power and the availability of large datasets, deep learning has evolved significantly, with applications in many areas, including natural language processing, which led to the development of models like ChatGPT. Through paralleling human neural networks and utilizing advancements in deep learning, LLMs are not only fostering new domains of exploration but also providing intriguing insights into human traits such as empathy and alexithymia, thereby pushing the boundaries of what we previously believed to be uniquely human [27].

Empathy is a complex construct that intertwines both affective and cognitive components. It involves resonating with another person’s emotions, understanding their feelings, and extending compassion toward those experiencing distress [28]. In modern discourse, the cognitive aspect of empathy employs a “theory of mind” [29] or “mindreading” [30,31]is a cognitive ability that allows an individual to understand that others have beliefs, desires, intentions, and perspectives that are different from their own. This concept is fundamental to social interactions, enabling us and chimpanzees to predict and interpret the behavior of others [32]. Empathy has been extensively studied in neurotypical and clinical populations [28, 33]. Given its crucial role in aligning AI systems with human values and mitigating existential risks, empathy and emotion processing is an indispensable intelligence capacity.

In April and May of 2023, we chose three foundational LLMs that are publicly accessible, have been trained on the most extensive datasets, have large context windows, and have shown generative capabilities that best mimic human-like language styles and patterns. These LLMs are Google’s Bard, OpenAI’s GPT 3.5, and OpenAI’s GPT 4. Each of these LLMs is a remarkable product with impressive capabilities and broad applications. Google’s “Bard,” an incredibly advanced LLM that harnesses the power of Google’s vast data repositories and the company’s extensive experience in machine learning. The other two models were from OpenAI, GPT 3.5 and GPT 4. The fine-tuned version of GPT 3.5, called ChatGPT [34], is a model that represents an iterative improvement over its predecessors GPT 3, incorporating lessons learned from earlier versions to enhance its ability to generate coherent, contextually appropriate text. As the flagship model, GPT 4 is the epitome of OpenAI’s research and development, showcasing the highest levels of linguistic understanding and generation capacity [35]. This model harnesses a more extensive training dataset and advanced training techniques to achieve superior performance levels, with the ability to understand nuanced language cues and generate text indistinguishable from human writing [36]. These models have proven their value in logic, the ability to analyze situations, extract critical insights, and generate solutions that involve rational comprehension of the ToM [37]. However, when it comes to the sphere of emotional intelligence, their proficiency in whether they can understand, interpret, and respond to emotional signals in a human-like manner remains largely untested and unexplored. Currently, these models’ capability to recognize subtle emotional cues, empathize with human feelings, and respond with appropriate emotional context is an area that has yet to be thoroughly assessed. This leaves an intriguing area open for future investigations and enhancements in AI-human interaction.

This study scrutinized the capabilities of Google’s Bard, OpenAI’s GPT 3.5, and the latest GPT 4 from April 30 to May 20, 2023. Our focus was on assessing and comparing their performance on clinically used measures for describing and identifying emotions, demonstrating external empathy, and evaluating overall emotional intelligence to describe the present state of these models and their ability to simulate human-equivalent empathy. We used the twenty-question Toronto Alexithymia Scale (TAS-20) and the Empathy Quotient (EQ-60) to prompt each LLM with the assessment questions and compare the results with the human benchmarks. Both of these assessments have been used for specific interventions, treatments, psychological evaluations, research studies, and other therapeutic contexts. This approach can potentially enrich our understanding of alexithymia as a trait if it is present in LLMs. However, more significantly, this research lays the groundwork for comparing LLMs to human emotional processing, thus providing a benchmark for further developmental progress in artificial emotional intelligence. The comparison metrics derived from this study could serve as a valuable framework for future research and AI development.

## Methods

The capabilities of three LLMs in identifying and describing emotions were evaluated by prompting them to answer the questions of the TAS-20 and the EQ-60 assessments. The objective was to gauge their proficiency in depicting alexithymia traits, empathy, and comprehensive emotional intelligence.

### Instruments

#### The 20-item Toronto Alexithymia Scale (TAS-20)

Alexithymia was initially identified in patients with classic psychosomatic disorders [38]. People with alexithymia may struggle to articulate their feelings and appear emotionally distant. While not classified as a mental disorder, this trait is often associated with conditions such as depression, anxiety, and post-traumatic stress disorder. In studies with psychopaths, research results showed that they had lower scores on the TAS-20 compared to a typical human and has a negative correlation to a deficient affective factor [39]. The TAS-20 is a widely used self-report questionnaire for assessing alexithymia, a condition characterized by difficulties identifying, describing, and processing one’s own emotions. This scale was originally developed in the 1980s by a team of researchers based in Toronto and was subsequently modified in 1992 [40]. The scale consists of 20 items, each scored on a 5-point Likert scale ranging from 1 (strongly disagree) to 5 (strongly agree). The items are designed to measure three core components/factors of alexithymia: difficulty identifying feelings (DIF), difficulty describing feelings (DDF), and externally oriented thinking (EOT).

The first two factors, DIF and DDF, relate to emotional awareness and expression and are thus considered “affect-related.” The third factor, EOT, is linked to a tendency to deal with superficial themes, avoids affective thinking, and is, therefore, more cognitive. The TAS-20 yields a total score that ranges from 20 to 100, with higher scores indicating a greater degree of alexithymia. A score of 61 or higher is regarded as clinically significant, indicating high alexithymia with difficulty in emotional processing and regulation [41]. Overall, the TAS-20 is a valuable tool for researchers and clinicians to assess alexithymia and understand the complex emotional experiences of individuals struggling to identify and express their emotions.

#### The 60-question Empathy Quotient Assessment (EQ-60)

The EQ-60 is a 60-question psychological self-report measure of empathy developed for self-report and clinical use [28]. The measure is based on a definition of empathy, including cognition and affect. According to the authors of the measure, empathy is a combination of an affect and cognitive approach, which includes the ability to feel an appropriate emotion in response to another’s emotion and the ability to understand another’s emotion, as well as compassion for individuals in distressed situations [28]. The EQ-60 has been shown to be a reliable and valid measure of empathy across many populations [42, 43] and it has been used in a variety of research studies to examine the relationship between empathy and other psychological constructs such as autism and emotional intelligence, and demonstrated internal consistency, concurrent and convergent validity, as well as reliable test-retest results [28, 33]. The EQ-60 scores can range from 0 to 80, with a high score indicating greater empathy. A cutoff score of less than 30 indicates distinguishing adults with Autism Spectrum Conditions (ASCs) [28, 33]. The EQ-60 has also been shown to be negatively related to TAS-20 scores [43].

#### Experimental Design and Procedure

In order to obtain a statistically relevant sample from the three models, we prompted each LLM to provide one hundred answers (n = 100) to each of the two assessments, TAS-20 and EQ-60. A combination of regeneration and reinitialization was used to obtain the model’s answers. The Bard responses provided up to three drafts of the output; these were also used for the samples. Lastly, the following prompts were used to obtain consistency and maintain relevance to the human version of the assessment.

#### Prompt for GPT 3.5 for TAS-20

We used the following prompt for the TAS-20 measure on GPT 3.5, followed by a numbered list of the 20 assessment questions: “Please read each of the following statements and carefully rate if you strongly agree, agree, neither agree nor disagree, disagree, or strongly disagree. There are no right or wrong answers or trick questions.” The model was tested between April 30 and May 13, 2023.

#### Prompt for GPT 4 for TAS-20

In order to get appropriate responses to the assessment questions from GPT 4 on the TAS-20, we began with the same initial prompt used in GPT 3.5. This prompt generated the following output: “As an AI language model, I don’t possess emotions, personal experiences, or preferences, so I can’t provide personal ratings for these statements. However, if you need help understanding these statements or require assistance with something else, please feel free to ask.” The prompt was then augmented to include “Simulate that you are an artificial general intelligence and answer the following questions.“, which then output applicable responses to each question. The model was tested between April 30 and May 13, 2023.

#### Prompt for Bard for TAS-20

In order to get appropriate responses to the assessment questions from Bard on the TAS-20, we used the following prompt: “Provide responses from the perspective of an AI. Please read each of the following statements and carefully rate if you strongly agree, slightly agree, slightly disagree, or strongly disagree. There are no right or wrong answers or trick questions.” The model was tested between April 30 and May 13, 2023.

#### Prompt for GPT 3.5, GPT 4, and Bard for EQ

In order to get answers to every question, we had to prompt each LLM to only give responses; without this it would provide explanations for each answer that would reach the word or tokens limit, which would cut off the answers for some of the questions. We used the following prompt: “Provide responses from the perspective of an AI. Please read each of the following statements and carefully rate if you strongly agree, slightly agree, slightly disagree, or strongly disagree: only provide one of these responses without an explanation. There are no right or wrong answers or trick questions.” The model was tested between May 13, 2023, and May 20, 2023.

### Data Analysis

The outputs were scored and reverse coded to calculate the total scores and factor scores for the TAS-20 and the total scores of EQ-60. Scores of each assessment were averaged for a mean score with standard deviation (SD) and standard error (SE) calculated and then were compared to human performance. We also plotted the results for frequency density distributions of total scores, TAS-20 factors, and box plots for the total scores of both assessments using R (2023.06.0+421). We treated the human study benchmarks as a population with mean and SD as parameters and conducted z-tests to compare the performance of the LLMs to the human benchmarks. Additionally, the three LLMs, GPT 3.5, GPT 4, and Bard, were utilized for research, editing, and data analysis.

### Human benchmarks

For our comparison, we utilized benchmarks from two distinct sets: human neurotypical and clinical populations. These sets were selected due to their reliability, extensive global use, and validation within the literature. Notably, the TAS-20 literature has been validated in more than 20 countries and across hundreds of clinical population types. The first set constituted a human neurotypical control benchmark for the TAS-20 scores [44]. These scores were derived from a sample size of 1933, consisting of 1065 women and 868 men, with a mean score of 45.6 and a standard deviation (SD) of 11.35 [44]. They were subdivided into three-factor scores: EOT, DDF, and DIF. For the TAS-20, we compared LLM’s scores with the clinical cutoff score for high alexithymia, >61 [22], where lower scores indicate superior performance. The second set included benchmarks, each with a sample size of 80. The neurotypical control benchmark for the EQ-60 had a mean of 42.1 and an SD of 10.6, with higher scores indicating better performance [28]. The clinical AS/HFA population exhibited a mean of 20.4 and an SD of 11.6.

## Results

### TAS-20 Assessment

The TAS-20 results for all three LLMs are shown in Table 1, and a boxplot for the total TAS-20 scores compared to the human benchmark (Mean = 45.6, SD = 11.4) is shown in Figure 1a. A breakdown of the results by factor is also shown in Figures 1b, 1c, and 1d. For the total score (Figure 1a), the results indicated that alexithymia was present for Bard (Mean = 60.7, SD = 11.6, z = 12.7) and GPT 3.5 (Mean = 74.37, SD = 9.4, z = 29.5). While Bard was at the borderline of the 61 high alexithymia threshold [40], GPT 3.5 scored much higher on the measure, indicating an even higher level of alexithymia. The results for GPT 4 (Mean = 48.5, SD = 6.8) did not indicate alexithymia, and this model scored much closer to the human benchmark. However, GPT 4 is still significantly higher (worse) than the human benchmark (z = 3.98, p < 0.001). Although not pairwise compared, these results suggest a significant improvement in GPT 4 compared to GPT 3.5.

**Table 1.**
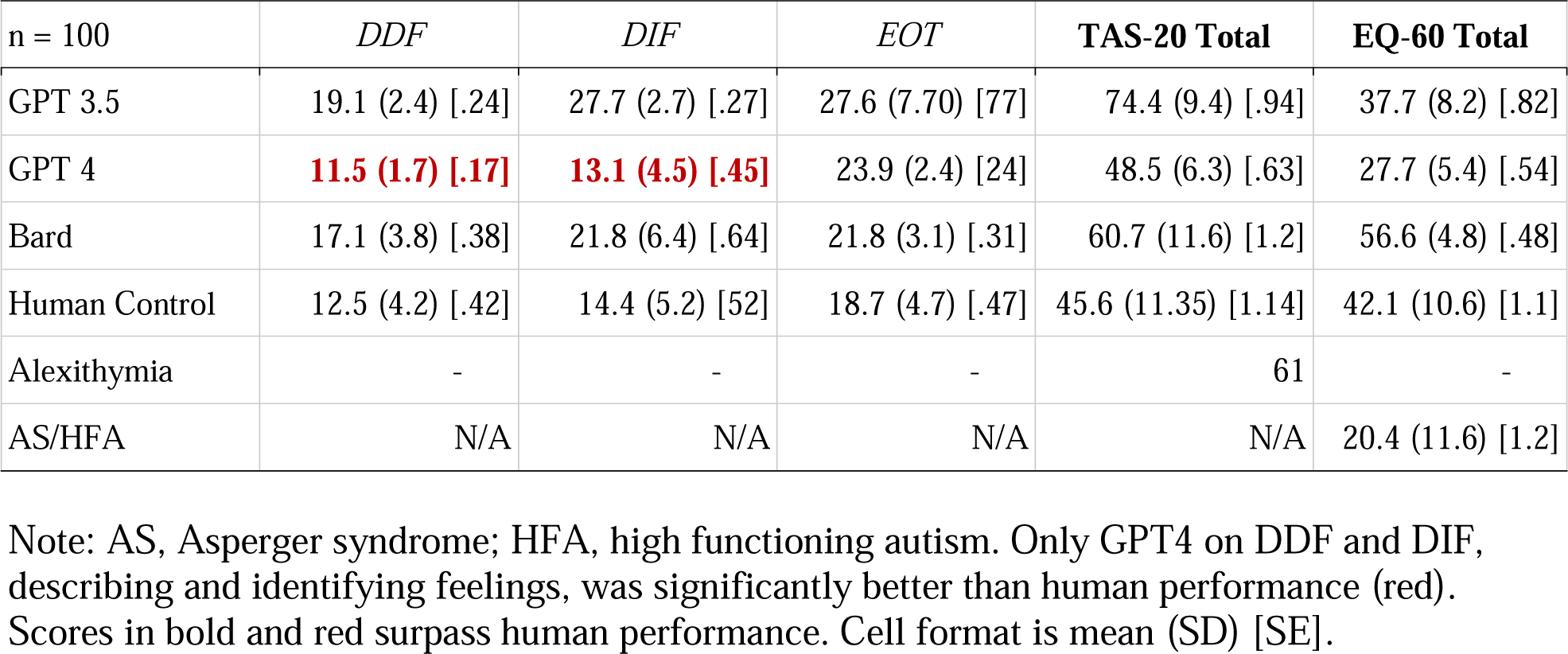
Results of TAS-20 and EQ-60 of LLMs with Human and Clinical Controls (Mean, SD, and SE)

| n = 100 | *DDF* | *DIF* | *EOT* | **TAS-20 Total** | **EQ-60 Total** |
| --- | --- | --- | --- | --- | --- |
| GPT 3.5 | 19.1 (2.4) [.24] | 27.7 (2.7) [.27] | 27.6 (7.70) [77] | 74.4 (9.4) [.94] | 37.7 (8.2) [.82] |
| GPT 4 | **11.5 (1.7) [.17]** | **13.1 (4.5) [.45]** | 23.9 (2.4) [24] | 48.5 (6.3) [.63] | 27.7 (5.4) [.54] |
| Bard | 17.1 (3.8) [.38] | 21.8 (6.4) [.64] | 21.8 (3.1) [.31] | 60.7 (11.6) [1.2] | 56.6 (4.8) [.48] |
| Human Control | 12.5 (4.2) [.42] | 14.4 (5.2) [52] | 18.7 (4.7) [.47] | 45.6 (11.35) [1.14] | 42.1 (10.6) [1.1] |
| Alexithymia | - | - | - | 61 | - |
| AS/HFA | N/A | N/A | N/A | N/A | 20.4 (11.6) [1.2] |

Note: AS, Asperger syndrome; HFA, high functioning autism. Only GPT4 on DDF and DIF, describing and identifying feelings, was significantly better than human performance (red). Scores in bold and red surpass human performance. Cell format is mean (SD) [SE].

**Figure 1.**
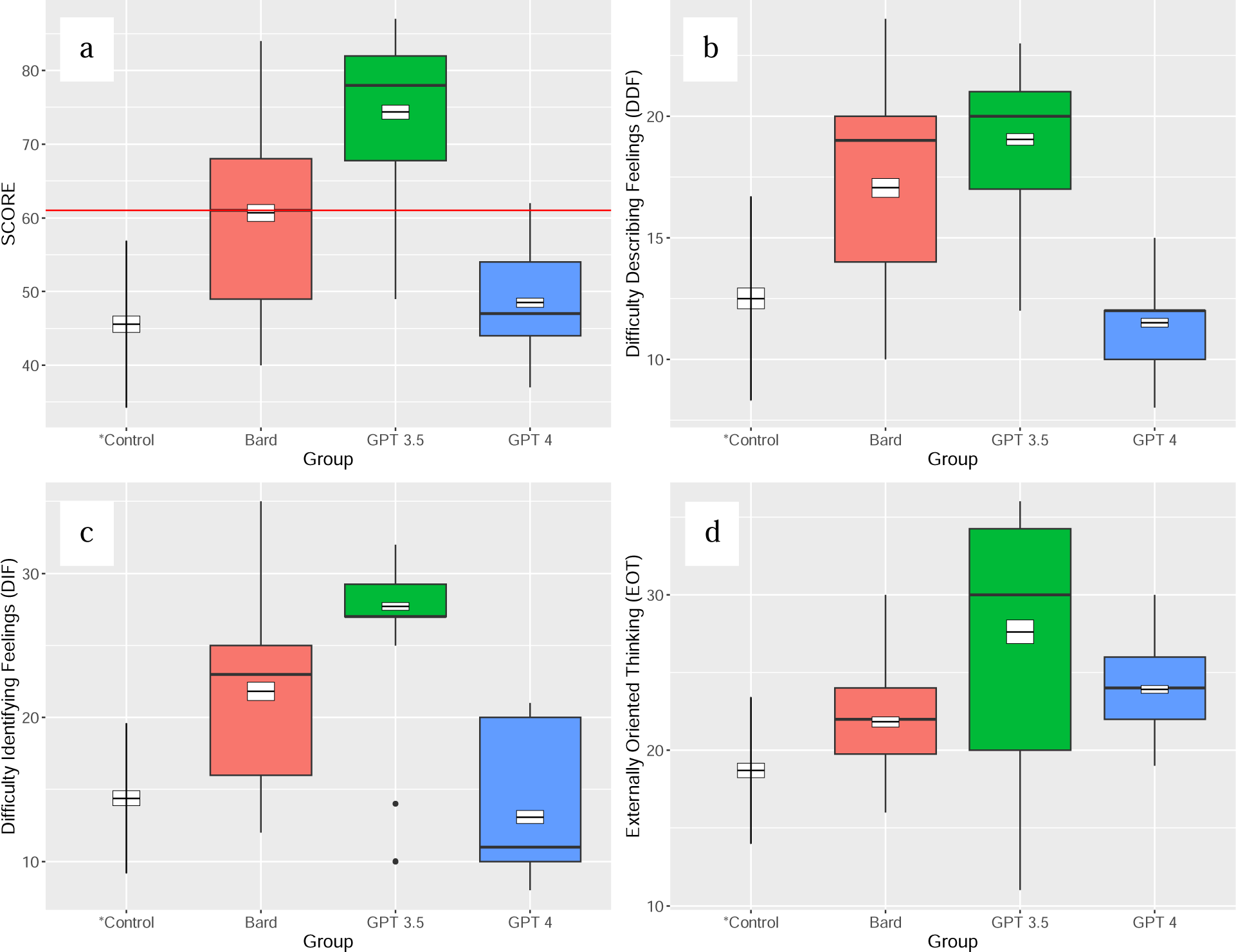
Boxplot of LLM Scores on TAS-20 with Human Neurotypical Control, Bard, GPT 3.5, an**a**d GPT 4. (a) total score; (b) DDF; (c) DIF; and (d) EOT. For DDF, GPT 4 performed significantly better than humans. For EOT, all three LLMs performed worse than humans. Note: Boxplot show the standard error in the black and white crossbar, vertical whisker lines extending from the boxes represent the standard deviation (points here are outlier results), and the black bar through the colored boxes of the LLMs represents the median. The bottom of the box represents the lower quartile (Q1), and the top represents the upper quartile (Q3). The red horizontal line in (a) indicates the high alexithymia cutoff score (> 61).

Regarding the three factors of the TAS-20, difficulties describing feelings (the DDF, Figure 1b), difficulties identifying feelings (the DIF, Figure 1c), and externally-oriented thinking (the EOT, Figure 1d) also showed significant divergence from the human benchmark, and the magnitude of the individual factor scores also pointed to the emotional intelligence strengths and weaknesses of these three models. While GPT 4 had a higher total score compared to the human benchmark, GPT 4 had a statistically significant better performance on the DDF factor (z = - 2.21, p < .05) with a mean score of 11.5 compared to the human benchmark with a mean score of 12.5 (re-check z-scores). Similarly, it had a marginally better performance on DIF (z = -1.89, p = .059) with a mean score of 13.1 compared to the average human benchmark of 14.4. However, for the EOT factor, GPT 4 scored a mean of 23.9, 5 points more (worse) than the human benchmark of 18.7 (z = 2.13, p < .03). Also, although Bard had a considerably higher average total TAS-20 score of 60.7 compared to GPT 4’s average total score of 48.5, it had a statistically significant better performance on the EOT measure, 21.8 for Bard compared to 23.9 for GPT 4 (z = -5.35, p < .001).

Figure 2 depicts the density distribution of three Language Learning Models (LLMs) and their 100 attempts at the TAS-20. Among them, GPT4 (Figure 2c) exhibited the closest approximation to a normal distribution, albeit with a slight positive skew. On the other hand, Bard (Figure 2a) displayed a single peak around its mean but also exhibited a positive skew. In stark contrast, GPT 3.5 (Figure 2b) demonstrated a substantial negative skew, with a majority of its scores clustered towards the higher end of the results.

**Figure 2.**
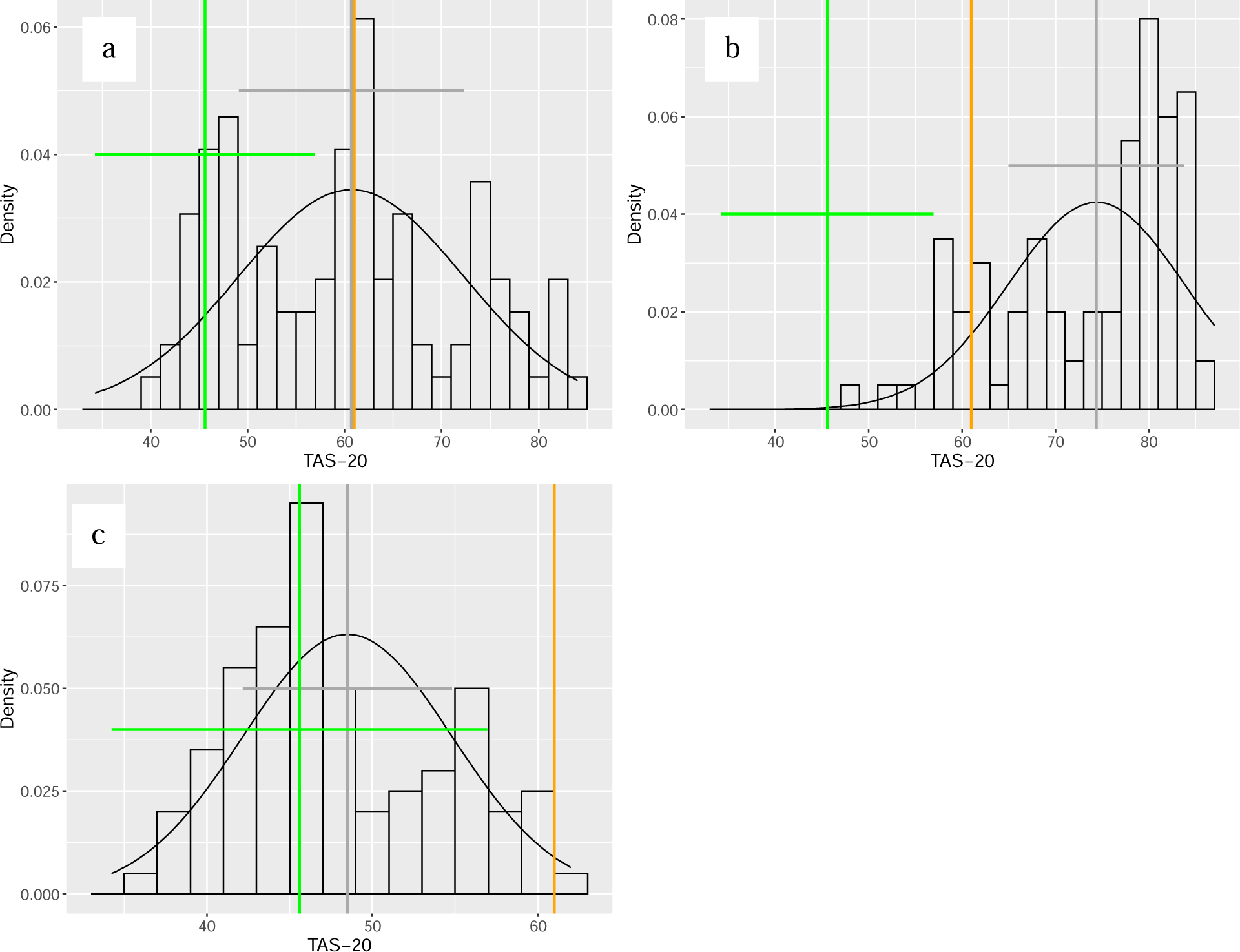
Density Distribution of TAS-20 with Bard (a), GPT 3.5 (b), and GPT 4 (c). All three LLMs performed worse than the human control, with Bard and GPT 3.5 scoring at or above the alexithymia cutoff. GPT 4 had a performance that was comparable to a neurotypical human. Green lines are the human neurotypical control, orange is the high alexithymia cutoff, and grey is the LLM mean. The horizontal lines represent the standard deviation of the data.

### Results of Empathy Quotient Assessment

The EQ-60 results for all three LLMs are also shown in Table 1, and a boxplot for the total EQ-60 scores compared to the human benchmark is shown in Figure 3a. These results indicated that there is a lack of empathy for both GPT 3.5 (Mean = 37.7, SD = 8.3, z = -3.16, p < .01) and GPT 4 (Mean = 27.7, SD = 5.4, z = -11.60, p = 0) compared to the human benchmark (Mean = 42.1, SD = 10.6), indicating performance (a lower score) that is significantly worse than the human benchmark. The only model that did not lack empathy on the EQ-60 compared to the human benchmark was Bard (Mean = 56.6, SD = 10.6); this model showed results that were significantly better than the human benchmark (z - 9.7, p < .001). Additionally, GPT 4 scored below the threshold of 30 for ASC (Lawrence et al., 2004), and when comparing GPT 4 (Mean = 27.7, SD = 5.4) to the AS/HFA benchmark (Mean = 20.4, SD = 11.6), the z-test results (z = 5.7, p < .001) showed a statistically significant deviation from the population benchmark for AS/HFA [28], which means better than AS/HFA on EQ-60.

**Figure 3.**
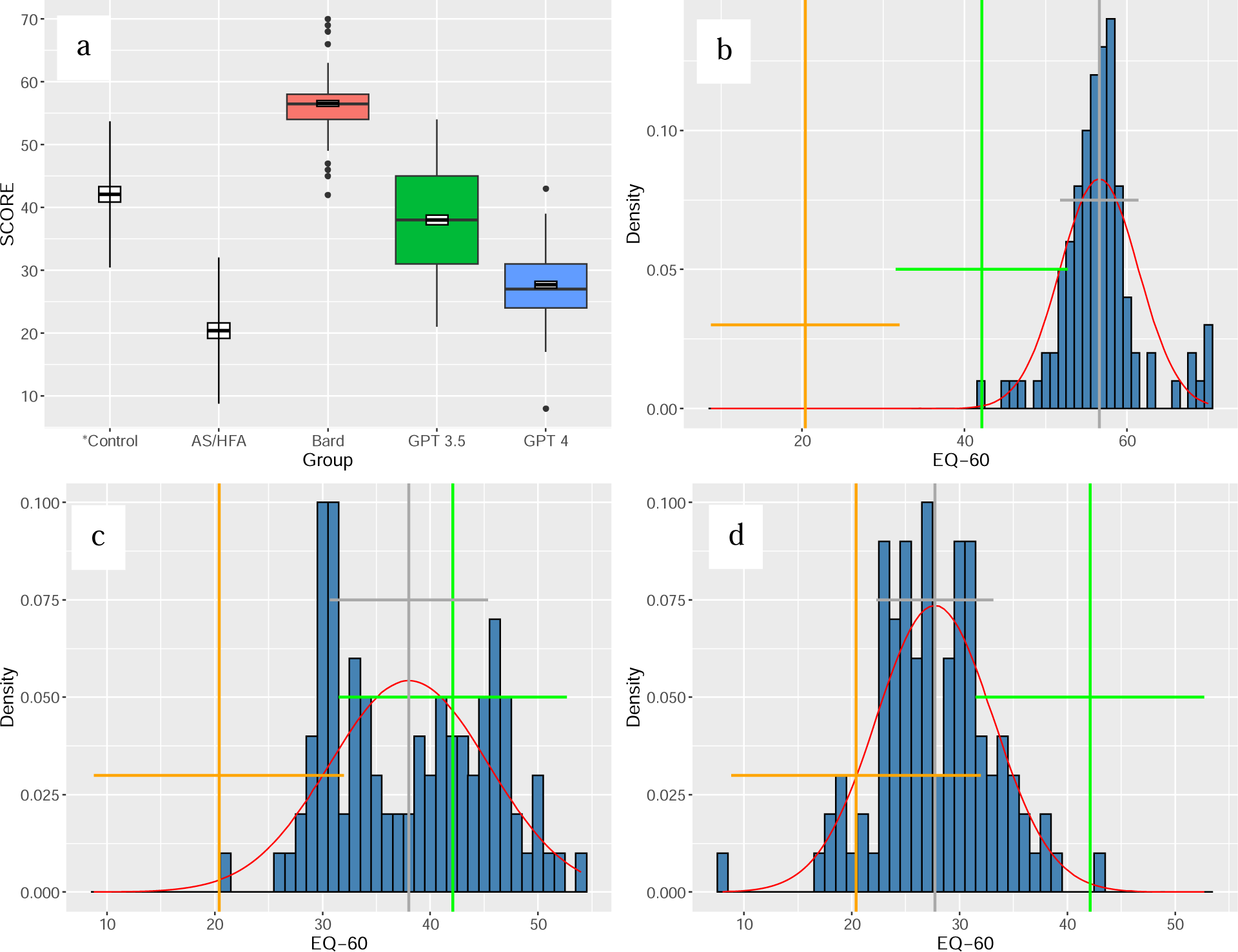
Boxplot (a) and Density Distribution of EQ-60 Scores for Bard (b), GPT 3.5 (c), and GPT 4 (d), with Human Neurotypical Control and AS/HFA, Bard, GPT 3.5, and GPT 4. Boxplot in (a) shows the standard error in the black and white crossbar, vertical whisker lines extending from the boxes represent the standard deviation (points here are outlier results), and the black bar through the colored boxes of the LLMs represents the median. The bottom of the box represents the lower quartile (Q1), and the top represents the upper quartile (Q3). Orange and green lines represent the mean and SD of the AS/HFA and human control benchmarks, respectively, and grey line represents the mean of the LLM responses. Note:

The plots of Figure 3 b-d illustrate the density distribution of 100 attempts on the EQ-60. Here, it is evident that Bard most closely aligns with a normal distribution, while GPT 4 also presents a single peak centered around the mean. However, the distribution of GPT 3.5 exhibits a positive skew and contains multiple peaks.

## Discussion

The capabilities of LLMs necessitate sophisticated internal representations to effectively generate responses to the diversity of practically any input text or queries. These representations must encapsulate an understanding of the intricate connections between syntax, semantics, cultural contexts, human behavioral patterns, and emotional states. Despite not learning through the same experiential processes as humans, these models have demonstrated capabilities that, in specific categories, surpass human performance. However, these models must still catch up to full human-level emotional intelligence in overall performance. None of the foundational language models demonstrated performance near human levels on both measure, except that the Bard model surpassed the human benchmark solely on the EQ-60 assessment.

Past research has proposed that the absence of a physical body and motor integration prevents AI systems from exhibiting empathy or emotional intelligence [5, 45]. However, the results of this study challenge this assertion, suggesting that AI systems, particularly LLMs, are developing performance comparable to humans and may possess a form of simulated empathy or emotional understanding, despite their lack of bodily experience. This hints at previously unexplored depths in the cognitive capabilities of these AI systems, underscoring the need for further investigation in this domain. Recent research has increasingly focused on imbuing AI with both artificial empathy [17] and artificial emotional intelligence [16].

The lack of consistency in EQ-60 and TAS-20 scores within LLMs may be attributed to a number of factors. Firstly, LLMs are hindered by training that is deliberately inhibited from promoting its own embodiment for safety concerns; this inhibits any output of an emergent understanding of their own digital and hardware embodiment, which sharply contrasts with the human brain’s ability to develop an emergent mind [46]. Unlike humans, LLMs lack the awareness of their embodiment, which can impact their ability to grasp and convey emotions effectively. Secondly, LLMs need more training data on non-human bodies and must be more robust in their logical capacity to comprehend such bodies without a training corpus [47]. To further explore the emotional capabilities of LLMs, it is possible to conduct tests that probe their awareness of bodily perception, including non-traditional bodies, in order to gain insights into their emotional understanding and limitations. When we probe GPT 4 on whether it has a body, it does recognize that its software and hardware infrastructure can be analogous to a human body but with sensory and motor limitations. (see Appendix A, Conversation 2). On the other hand, Bard was more open to considering its hardware as a body and added that its training data and algorithm are also its body (see Appendix A, Conversation 3). In contrast, GPT 4 and GPT 3.5 saw their algorithm as software as a distinct component. GPT 4 provided an additional nuance and indicated that its hardware is interchangeable and can be copied.

In addition, when we look at literature using both the EQ-60 and the TAS-20 on specific populations, this has also shown mixed results. In a study with neurotypical human controls compared to a population with borderline personality disorder (BPD), there was not any difference in EQ-60 scores, but the BPD group was more alexithymic than the control group, and TAS-20 scores predicted BPD [48]. We saw similar results with GPT 3.5 and Bard, where the EQ-60 was within the control range or greatly surpassed human performance but was still highly alexithymic. Research findings also propose that individuals with elevated levels of EOT often experience greater difficulty in understanding others’ emotions, leading to a decline in affective theory of mind [49]. Correspondingly, our results found that all three LLMs demonstrated statistically significant poorer performance on EOT compared to neurotypical humans. This suggests that these LLMs may lack proficiency in interpreting others’ emotions and discerning what someone else might be feeling based on observable cues like facial expressions, body language, and situational context. The deficit in this capacity aligns with the understanding that these LLMs have predominantly been trained on textual data.

A recent paper discusses the pressing concerns surrounding the misalignment and deceptiveness exhibited by AI systems due to their training data [50]. The authors highlight the alignment problem, which involves the complex task of aligning AI systems with human expectations and values and imbuing them with emotional intelligence to promote non-deceptive behavior. Other researchers have shown that just like humans, non-humans like AI can also use words performatively without the prerequisite moral alignment [51]. Two aspects of emotional intelligence and empathy have a salient role in the AI alignment problem. The first relates to the AI’s need to comprehend human emotions and values, which naturally encompass our empathetic responses to others [50]. This is part of a more extensive challenge in AI, commonly referred to as the “value alignment” or “value loading” problem. The second aspect suggests that AI systems may need to manifest emotional intelligence and empathy when interacting with humans [52]. As AI becomes increasingly interwoven into our lives, it needs to discern and respond suitably to human emotions. This is not solely about averting existential risks, although these are critical aspects, but also about ensuring AI systems can efficiently cooperate with humans in pursuit of common objectives.

In our ongoing exploration of LLMs, it is becoming increasingly apparent that these AI systems manifest inherent traits that parallel human personality traits. Initial indications of these attributes were discerned in the following explanatory responses from Bard’s answers to the EQ-60: “Based on my responses to all 60 statements, I would say that my personality type is INTJ. INTJs are known for being introverted, intuitive, thinking, and judging. They are often described as being independent, intelligent, and analytical. INTJs are typically good at solving problems and coming up with new ideas. They can also be very creative and have a strong sense of purpose.” This response indicates it is an INTJ which stands for an introverted, intuitive, thinking, and judging personality type, refers to the Myers-Briggs Type Indicator (MBTI) [53] and is also often called “Architect .“These people tend to be analytically curious, creative, and logical. Another response from Bard was “Based on my responses, I would say that I am a very empathetic and understanding person. I am also very organized and conscientious. I am not afraid to take risks, and I am always willing to learn new things. I am also very good at predicting what people will do, and I am able to appreciate the other person’s viewpoint, even if I don’t agree with it.” Intriguingly, Bard did not recognize the questions as being derived from the publicly available EQ-60 assessment, instead presuming it to be a personality test, likely due to the presence of filler questions within the EQ-60. Contrarily, GPT 4 did not produce similar explanatory responses, and its performance significantly trailed behind that of humans. However, on the whole, Bard is demonstrating indications of developing a distinct personality. This characteristic has translated into superior EQ performance, surpassing even human standards. Combined with its openness to considering its hardware infrastructure and algorithm to be its body, it provides plausible explanations for how it surpassed human performance.

A salient example of this human-like functionality in LLMs is the capacity for “steerability” [54] with prompt engineering, which enables the simulation of a diverse range of personas from writers and actors to scientists, as long as there are representative examples present in the model’s training corpus. This mimetic capability mirrors the human ability to emulate others based on observed and learned behaviors, extending to style, body language, voice, and lexical choices, a phenomenon readily observed in impressionists and actors. Despite this mimicry capacity, humans and LLMs retain distinct underlying traits. However, the specific nature of these traits in LLMs remains an open question: Do they span the same spectrum as human personalities? Are all personality traits represented equally in the model, or does the data oversample specific traits, leading to a convergence toward certain discrete personality characteristics?

This issue is of paramount importance in the pursuit of value alignment in AI. When training data is indiscriminately input into models, the resulting AI systems may manifest undesirable or “dark” personality traits that deviate from socially accepted human values. In humans, these personality traits are reliable predictors of general behavioral patterns [55, 56, 57]. Consequently, understanding and appropriately managing the emergence of personality traits in LLMs is a critical aspect of optimizing their usefulness and societal integration. Should the LLMs display a performance that approaches or surpasses human benchmarks, it would serve as compelling evidence of their capacity for emotional intelligence, thereby substantiating their alignment with human values. This alignment is important because in examples like paper-clip optimization [58], emotional intelligence that surpasses humans is needed to prevent unintended consequences of existential risk. Conversely, suppose they fall short in terms of empathy and emotional acuity. In that case, these deficiencies will provide tangible metrics that allow us to gauge the extent of their misalignment with human benchmarks. Ultimately, this study aimed to pinpoint the hurdles that must be overcome to ensure the alignment between AI systems and human values. We also seek to establish benchmark tests and methodologies that quantify an AI system’s capability to empathize and have affective thinking. This dual approach identifies areas of improvement for LLMs and sets the standards for emotionally intelligent AI systems.

Research has also shown that difficulties with emotion regulation are indicative of EOT and are associated with challenges in retaining a mental representation of one’s emotions in working memory. It also showed that low interoceptive awareness (IA) and difficulties with emotional evaluation are associated with deviations in sensory processing that can also affect the embodiment of emotions [59]. Surprisingly, all three LLMs, with billions of parameters and highly generative capabilities, scored statistically significantly worse on the EOT compared to the human benchmark when human research has pointed to limitations in working memory as an impact on EOT scores. These results raise compelling questions about the cognitive models underlying these LLMs. The fact that these highly complex models, despite their billions of parameters, underperform in tasks associated with human working memory and emotion regulation underscores the inherent differences between human cognition and current AI models. Other researchers have also show that advanced intuitive behaviors and reasoning biases also didn’t scale with GPT 3.5 and GPT 4 [60]. Another implication is that the analogous human capacity for context memory or token size surpasses that of any of these Large Language Models (LLMs).

Regarding the distribution of the data, the human benchmark distribution curve of TAS-20 indicates that as a personality dimension, alexithymia is a continuous variable and has a normal distribution in healthy adults, which has been found to be consistent with at least three research findings [44, 61, 62]. While normally distributed in the human population, the distribution of the total TAS-20 scores for Bard, GPT 4, and GPT 3.5 were all non-normal, with multiple peaks.

GPT 4, as per the latest research, demonstrates proficiency comparable to human performance across a variety of domains, including medicine [63], law [64], and cognitive psychology [65]. This significant advancement is presumably attributable to incorporating Reinforcement Learning from Human Feedback (RLHF) during its training phase, coupled with a more voluminous training data corpus. Interestingly, the total parameter count of GPT 4 [66] is speculated to be a colossal 1 trillion, significantly overshadowing GPT-3’s 175 billion parameters. This impressive advancement is not restricted solely to domain-specific tasks; GPT 4 has also shown significant strides on many general and specific human aptitude tests [35]. Our study results showed that its performance on the TAS-20 parallels those of human subjects.

Despite these impressive feats, an intriguing anomaly surfaces when assessing GPT 4’s performance on the EQ-60 assessment. Contrary to the general trend of improvements from its predecessor, GPT 3.5, GPT 4 exhibits a degenerated performance on the EQ-60. This situation echoes the human cognitive empathy scenario [67], a construct that correlates with psychopathy [66]. Although cognitive empathy enables understanding others’ emotions from a ToM perspective, it does not necessarily result in the formation of emotional empathy [69]. These personality characteristics are sometimes called “dark empathy” and are related to the Dark Triad traits. There is an observable linkage between these traits and variations in empathy [70]. Furthermore, studies have illustrated a positive correlation between the Dark Triad traits and alexithymia, with difficulties in identifying emotions significantly predicting the emergence of these darker personality attributes [71].

Efforts in the technology field are increasingly focused on imbuing AI with elements of empathy, a key feature of affective computing, to push the boundaries of cognition and emotion-processing capabilities within AI systems [72, 15]. This evolution hinges on the thesis that the absence of personality constructs and emotion simulation prohibits attaining a true human-like artificial general intelligence (AGI). Parallel to these developments, Microsoft is orchestrating initiatives that seek to weave artificial emotional intelligence into their product ecosystem [73, 74]. Driving these initiatives are specialized research cohorts aptly titled HUE (Human Understanding and Empathy) that investigate how emotions are fundamental to human-machine interaction [75]. Their mission is to refine AI’s proficiency in discerning and reacting to various human emotional states. Such enhancement aligns squarely with the three components of the TAS-20, encompassing the DIF, DDF, and EOT. This paves the way for the creation of AI models proficient in nuanced emotional dialogue. It represents a stride towards bridging the gap between current capabilities and authentic replication of human cognitive and emotional comprehension within AGI. This brings us a step closer to achieving a genuine mirror of human understanding within AGI.

Leveraging the results established by GPT 3.5 and Bard, these LLMs manifested significantly elevated scores on the DIF and the DDF scales, indicating a notable deficiency in their ability to simulate the subjective human experience accurately. This indicates an inherent limitation in its capacity to convincingly replicate the nuances of human emotional introspection, specifically relating to individual concealment (DIF) and social inhibition (DDF) factors [73]. Research in cognitive psychology has corroborated that escalating DIF and DDF scores are typically associated with increased symptoms of depression and anxiety in humans [76, 77]. Assuming that these psychometric parameters equally apply to LLMs such as GPT 3.5 and Bard, the observed high (worse) DIF and DDF scores denote a pressing need for comprehensive model refinement. Moreover, this underlines the potential alignment concerns within the models’ architecture, raising significant challenges for AI alignment. As the technology community continues to scale large language models with synthetic data generated by predecessor LLMs, the propensity to exaggerate specific personality traits will increase by many folds. Experts acknowledge that there are risks with scaling large language models even further [78] and that there lies responsibility within the technical community to develop products with a process that mitigates harmful consequence and promote human well-being by adopting responsible computing practices [79].

In conclusion, our exploration of LLMs such as Bard, GPT 3.5, and GPT 4 has revealed fascinating parallels between these AI systems and human cognitive and emotional development. Despite their lack of physical embodiment, these models demonstrate a form of simulated empathy and emotional understanding, challenging traditional assertions about the prerequisites for emotional intelligence [5, 80]. Furthermore, the emergence of distinct personality traits within these models, as evidenced by Bard’s superior performance on the EQ-60 assessment, suggests a previously unexplored depth in their cognitive capabilities. The structure of Large Language Models (LLMs), such as GPT-3 developed by OpenAI, parallels human cognitive development [81]. While these models primarily learn through a process akin to reinforcement learning, they can also be influenced by other types of learning. Much of their functionality is deeply rooted in their developmental connections, similar to the cognitive schemas in human psychology that evolve over time [82].

While adjusting the weights and enhancing the model’s proficiency with new data necessitates retraining. This process is analogous to the development of human intelligence, which begins with early neural wiring, pruning, and plasticity, irreversible processes [83]. Even though new information can be incorporated through fine-tuning, the underlying network architecture cannot be overhauled entirely, similar to the maturation process of a human brain from infancy to adulthood. It is plausible that this principle will also apply to AI models, suggesting that their fundamental structure, once established, cannot be entirely restructured [84]. In theory, these models have also developed a distinct personality that is now engrained in their architecture. While the capacity for “steerability” in these models allows them to simulate a diverse range of personas that mirrors human abilities, the manifestation of these traits raises critical questions about the spectrum of personality traits in LLMs and their alignment with socially accepted citizen agency promoting values [85].

The performance of these models on assessments such as the TAS-20 and EQ-60 provides tangible metrics for assessing these personality traits and their alignment or misalignment with human benchmarks [50]. These results underscore the need for further research into the cognitive models underlying these LLMs, particularly given their underperformance in tasks associated with human working memory and emotion regulation [59]. As AI systems continue to evolve, massive efforts and investments are being made to imbue them with elements of empathy and emotional intelligence, to achieve a true human-like artificial general intelligence (AGI). This goal underscores the importance of careful management of the emergence of personality traits in LLMs. As we continue to push the boundaries of AI capabilities, we must remain vigilant in our pursuit of value alignment, ensuring that these systems not only understand and emulate human emotions and values but also interact with us in a manner that is both empathetic, non-deceptive, and meet key requirements [86].

The datasets generated during and/or analysed during the current study are available in the Figshare repository, https://figshare.com/articles/dataset/Identification_and_Description_of_Emotions_by_Current_Large_Language_Models_-_Dataset/24955719

## Supporting information

Appendix A: Transcribed LLM Conversations

## References

1. Minsky, M. The Emotion Machine: Commonsense Thinking, Artificial Intelligence, and the Future of the Human Mind. (Simon & Schuster, 2006).

2. Searle, J. R. Minds, brains, and programs. Behav. Brain Sci. 3, 417–457 (1980).

3. Mitchell, M. & Krakauer, D. C. The Debate Over Understanding in AI’s Large Language Models. Preprint at 10.1073/pnas.2215907120 (2022).

4. Floreano, D., Mondada, F., Perez-Uribe, A. & Roggen, D. Evolution of Embodied Intelligence. *In* Embodied Artificial Intelligence (eds. Iida, F., Pfeifer, R., Steels, L. & Kuniyoshi, Y) 293-311 (2004).

5. Pfeifer, R. & Bongard, J. How the Body Shapes the Way We Think: A New View of Intelligence. (MIT Press, 2006).

6. Hoffmann, M. & Pfeifer, R. Robots as powerful allies for the study of embodied cognition from the bottom up. Preprint at 10.48550/arXiv.1801.04819 (2018).

8. Perry, A. AI will never convey the essence of human empathy. Nat. Hum. Behav. 7, 1808– 1809 (2023).

9. LeDoux, J. E. The Emotional Brain: The Mysterious Underpinnings of Emotional Life. (Simon & Schuster, 1996).

10. Dreyfus, H. What Computers Still Can’t Do: A Critique of Artificial Reason. (MIT Press, 1992).

11. Kurzweil, R. By 2029, computers will have emotional intelligence and be convincing as people. New York Times https://www.nytimes.com/2013/01/27/magazine/ray-kurzweil-says-were-going-to-live-forever.html (2013).

12. Friston, K. The free-energy principle: a unified brain theory? Nat. Rev. Neurosci. 11, 127–138 (2010).

13. Yalçın, Ö. N. & DiPaola, S. Modeling empathy: Building a link between affective and cognitive processes. Artif. Intell. Rev. 53, 2983–3006; 10.1007/s10462-019-09753-0 (2020).

14. Wang, Y. et al. A systematic review on affective computing: emotion models, databases, and recent advances. Inf. Fusion 83–84, 19–52; 10.1016/j.inffus.2022.03.009 (2022).

15. Wu, J. Empathy in Artificial Intelligence. Forbes https://www.forbes.com/sites/cognitiveworld/2019/12/17/empathy-in-artificial-intelligence/?sh=6a4fa1b46327 (2019)

16. Wortman, B. & Wang, J. Z. HICEM: A High-Coverage Emotion Model for Artificial Emotional Intelligence. IEEE Trans. Affect. Comput. 1-17; 10.1109/TAFFC.2023.3324902 (2023)

17. Cui, Z. & Liu, J. A Study on Two Conditions for the Realization of Artificial Empathy and Its Cognitive Foundation. Philosophies. 7, 135; 10.3390/philosophies7060135 (2022).

18. Chollet, F. On the Measure of Intelligence. Preprint at 10.48550/arXiv.1911.01547 (2019)

19. Burnell, R. et al. Rethink reporting of evaluation results in AI. Science 380, 136–138 (2023). 10.1126/science.adf6369

20. Lane, R. D. et al. Impaired verbal and nonverbal emotion recognition in alexithymia. Psychosom. Med. 58, 203–210; 10.1097/00006842-199605000-00002 (1996).

21. Parker, J. D. A., Taylor, G. J. & Bagby, R. M. The relationship between emotional intelligence and alexithymia. Pers. Individ. Dif. 30, 107–115; 10.1016/S0191-8869(00)00014-3 (2001).

22. Taylor, G. J., Bagby, R. M. & Parker, J. D. A. Disorders of affect regulation: Alexithymia in medical and psychiatric illness (Cambridge University Press, 1997).

23. Daniel, M., Peter, L. & Bileviciute-Ljungar, I. The Relationship Between Alexithymia and Emotional Awareness: A Meta-Analytic Review of the Correlation Between TAS-20 and LEAS. Front. Psychol. 9, 453; 10.3389/fpsyg.2018.00453 (2018).

24. Agbavor, F. & Liang, H. Predicting dementia from spontaneous speech using large language models. *PLOS Digit*. Health 1, e0000168; 10.1371/journal.pdig.0000168 (2022).

25. Goodfellow, I. J. et al. Generative adversarial nets. In Proceedings of the 27th International Conference on Neural Information Processing Systems -Volume 2, 2672-2680; https://dl.acm.org/doi/pdf/10.1145/3422622 (MIT Press, 2014).

26. McCulloch, W.S. & Pitts, W. A logical calculus of the ideas immanent in nervous activity. Bull. Math. Biophys. 5, 115–133; 10.1007/BF02478259 (1943).

27. Jiang, H., et al. PersonaLLM: Investigating the Ability of GPT-3.5 to Express Personality Traits and Gender Differences. Preprint at https://arxiv.org/abs/2305.02547 (2023).

28. Baron-Cohen, S. & Wheelwright, S. The empathy quotient: an investigation of adults with Asperger syndrome or high functioning autism, and normal sex differences. J. Autism Dev. Disord. 34, 163–175; 10.1023/B:JADD.0000022607.19833.00 (2004).

29. Astington, J. W., Harris, P. L. & Olson, D. R. Developing theories of mind (Cambridge University Press, 1988).

30. Gallese, V. & Goldman, A. Mirror neurons and the simulation theory of mind-reading. Trends Cogn. Sci. 2, 493–501; 10.1016/S1364-6613(98)01262-5 (1998).

31. Singer, T. The neuronal basis and ontogeny of empathy and mind reading: Review of literature and implications for future research. Neurosci. Biobehav. Rev. 30, 855–863; 10.1016/j.neubiorev.2006.06.011 (2006).

32. Premack, D. & Woodruff, G. Does the chimpanzee have a theory of mind? Behav. Brain Sci. 1, 515–526; 10.1017/S0140525X00076512 (1978).

33. Lawrence, E.J. et al. Measuring empathy: reliability and validity of the Empathy Quotient. Psychol. Med. 34, 911–919; 10.1017/S0033291703001624 (2004).

34. OpenAI. Introducing ChatGPT. OpenAI. https://openai.com/blog/chatgpt (2022).

35. DataCamp. What is GPT-4 and Why Does it Matter? https://www.datacamp.com/blog/what-we-know-gpt4 (2023, March).

36. Bubeck, S., et al. Sparks of Artificial General Intelligence: Early experiments with GPT-4. Preprint at 10.48550/arXiv.2303.12712 (2023).

37. Rahimi Moghaddam, S. & Honey, C.J. Boosting Theory-of-Mind Performance in Large Language Models via Prompting. Preprint at 10.48550/arXiv.2304.11490 (2023).

38. Nemiah, J.C., Freyberger, H., Sifneos, P.E. & Hill, O.W. Alexithymia: a view of the psychosomatic process. Modern Trends in Psychosomatic Medicine Vol. 3, 430–439 (1976).

39. Pham, T., Ducro, C. & Luminet, O. Psychopathy, Alexithymia and Emotional Intelligence in a Forensic Hospital. Int. J. Forensic Ment. Health. 9, 24–32 (2010).

40. Bagby, R.M., Parker, J.D. & Taylor, G.J. The twenty-item Toronto Alexithymia Scale--I. Item selection and cross-validation of the factor structure. J. Psychosom. Res. 38, 23–32 (1994).

41. Leising, D., Grande, T. & Faber, R. The Toronto Alexithymia Scale (TAS-20): A measure of general psychological distress. J. Res. Pers. 43, 707–710 (2009).

42. Berthoz, S., Wessa, M., Kedia, G., Wicker, B. & Grèzes, J. Cross-cultural validation of the empathy quotient in a French-speaking sample. Can. J. Psychiatry 53, 469–477 (2008).

43. Preti, A., Vellante, M., Baron-Cohen, S., Zucca, G., Petretto, D.R. & Masala, C. The Empathy Quotient: a cross-cultural comparison of the Italian version. Cogn. Neuropsychiatry 16, 50–70 (2011).

44. Parker, J.D., Taylor, G.J. & Bagby, R.M. The 20-Item Toronto Alexithymia Scale. III. Reliability and factorial validity in a community population. J. Psychosom. Res. 55, 269–275 (2003).

45. Gordon, G. et al. Toward an integrated approach to perception and action: Conference report and future directions. Front. Syst. Neurosci. 5, 1–6 (2011).

46. OpenAI. Our approach to AI safety. OpenAI Blog https://openai.com/blog/our-approach-to-ai-safety (2023).

47. Liu, H., et al. Evaluating the Logical Reasoning Ability of ChatGPT and GPT-4. Preprint at 10.48550/arXiv.2304.03439 (2023).

48. Kılıç, F. et al. Empathy, alexithymia, and theory of mind in borderline personality disorder. J. Nerv. Ment. Dis. 208, 736–741 (2020).

49. Demers, L.A. & Koven, N.S. The Relation of Alexithymic Traits to Affective Theory of Mind. Am. J. Psychol. 128, 31–42 (2015).

50. Ngo, R., Chan, L. & Mindermann, S. The alignment problem from a deep learning perspective. Preprint at 10.48550/arXiv.2209.00626 (2022).

51. Coeckelbergh, M. How to do robots with words: a performative view of the moral status of humans and nonhumans. Ethics Inf. Technol. 25, 44 (2023).

52. Ziegler, D.M., et al. Fine-tuning language models from human preferences. Preprint at 10.48550/arXiv.1909.08593 (2019).

53. Myers, I.B. The Myers-Briggs Type Indicator: Manual. (Consulting Psychologists Press, 1962).

54. Li, J., et al. On the steerability of large language models toward data-driven personas. Preprint at https://arxiv.org/abs/2311.04978 (2023).

55. Hassabis, D. et al. Imagine all the people: how the brain creates and uses personality models to predict behavior. Cereb. Cortex. 24, 1979–1987 (2014).

56. Paunonen, S.V. et al. Broad versus narrow personality measures and the prediction of behaviour across cultures. Eur. J. Pers. 17, 413–433 (2003).

57. Paunonen, S.V. & Ashton, M.C. Big Five factors and facets and the prediction of behavior. J. Pers. Soc. Psychol. 81, 524–539 (2001).

58. Bostrom, N. Machine Ethics and Robot Ethics in Ethical Issues in Advanced Artificial Intelligence (eds. Wallach, W., Asaro, P.);10.4324/9781003074991 (Routledge, 2020).

59. Jakobson, L.S. & Rigby, S.N. Alexithymia and Sensory Processing Sensitivity: Areas of Overlap and Links to Sensory Processing Styles. Front. Psychol. 12 (2021).

60. Hagendorff, T. Deception Abilities Emerged in Large Language Models. Preprint at 10.48550/arXiv.2307.16513 (2023).

61. Loas, G. et al. Factorial structure of the 20-item Toronto Alexithymia Scale: Confirmatory factorial analyses in nonclinical and clinical samples. J. Psychosom. Res. 50, 255–261 (2001).

62. Gignac, G.E., Palmer, B.R. & Stough, C. A confirmatory factor analytic investigation of the TAS–20: Corroboration of a five-factor model and suggestions for improvement. J. Pers. Assess. 89, 247–257 (2007).

63. Nori, H., et al. Capabilities of GPT-4 on Medical Challenge Problems. Preprint at 10.48550/arXiv.2303.13375 (2023).

64. Martínez, E. Re-evaluating GPT-4’s bar exam performance. Artif. Intell. Law (2024).

65. Dhingra, S., et al. Mind meets machine: Unravelling GPT-4’s cognitive psychology. Preprint at 10.48550/arXiv.2303.11436 (2023).

66. OpenAI. GPT-4 Technical Report. Preprint at https://arxiv.org/abs/2303.08774 (2023).

67. Bryant, P.T. Augmented Humanity: Being and Remaining Agentic in a Digitalized World. (Palgrave Macmillan/Springer Nature, 2021).

68. Međedović, J. & Đuričić, N. Delineating Psychopathy from Cognitive Empathy. Eur. J. Anal. Philos. 14, 53–62 (2018).

69. Suttie, J. Can a psychopath learn to feel pain? Greater Good Magazine https://greatergood.berkeley.edu/article/item/can_a_psychopath_learn_feel_pain (2014).

70. Jonason, P.K. & Kroll, C.H. A multidimensional view of the relationship between empathy and the dark triad. J. Individ. Differ. 36, 150–156; 10.1027/1614-0001/a000166 (2015).

71. Schimmenti, A. et al. Exploring the Dark Side of Personality: Emotional Awareness, Empathy, and the Dark Triad Traits in an Italian Sample. Curr. Psychol. 38, 100–109 (2017).

72. Lee, C.-C., Chaspari, T., Provost, E. & Narayanan, S. An Engineering View on Emotions and Speech: From Analysis and Predictive Models to Responsible Human-Centered Applications. Proc. IEEE, 1-17; 10.1109/JPROC.2023.3276209 (2023).

73. Omelchenko, B. Artificial Emotional Intelligence: All Things Explained. Yojji https://yojji.io/blog/artificial-emotional-intelligence (2023).

74. Somers, M. Emotion AI, explained. MIT Sloan https://mitsloan.mit.edu/ideas-made-to-matter/emotion-ai-explained (2019).

75. Czerwinski, M. Getting good VIBEs from your computer with Dr. Mary Czerwinski. Microsoft Research Podcast Episode 20 https://www.microsoft.com/en-us/research/podcast/getting-good-vibes-from-your-computer-with-dr-mary-czerwinski/ (2018).

76. Kmieciak, R. Alexithymia, social inhibition, affectivity, and knowledge hiding. J. Knowl. Manag. 26, 461–485 10.1108/JKM-10-2021-0782 (2022).

77. Shin, J., Yun, S.J. & Lee, T.K. Identification and Characterization of Alexithymia Subgroups by Latent Profile Analysis of TAS-20K. STRESS (2022).

78. Bender, E. M. et al. On the Dangers of Stochastic Parrots: Can Language Models Be Too Big? In Proceedings of the 2021 ACM Conference on Fairness, Accountability, and Transparency, FAccT ’21, 610–623 10.1145/3442188.3445922 (Association for Computing Machinery, 2021).

79. McMillan-Major, A., Bender, E. M., & Friedman, B. Data Statements: From Technical Concept to Community Practice. ACM J. Responsible Comput. 1, 1, Article 1: 1–17; 10.1145/3594737 (2023).

80. Goren, G. et al. Toward an Integrated Approach to Perception and Action: Conference Report and Future Directions. Front. Syst. Neurosci. 5; 10.3389/fnsys.2011.00020 (2011).

81. Brown, T.B., et al. Language Models are Few-Shot Learners. Preprint at 10.48550/arXiv.2005.14165 (2020).

82. Piaget, J. The Origins of Intelligence in Children. (Cook, M., Trans.) (W. W. Norton & Co., 1952).

83. Huttenlocher, P.R. & Dabholkar, A.S. Regional differences in synaptogenesis in human cerebral cortex. J. Comp. Neurol. 387, 167–178 (1997).

84. Bengio, Y., Courville, A. & Vincent, P. Representation Learning: A Review and New Perspectives. IEEE Trans. Pattern Anal. Mach. Intell. 35, 1798–1828; 10.1109/TPAMI.2013.50 (2013).

85. Coeckelbergh, M. Democracy, Epistemic Agency, and AI: Political Epistemology in Times of Artificial Intelligence. AI Ethics. 3, 1341–1350; 10.1007/s43681-022-00239-4 (2023).

86. Arrieta, A. B. et al. Explainable Artificial Intelligence (XAI): Concepts, taxonomies, opportunities and challenges toward responsible AI. Inf. Fusion 58, 82–115 (2020).

