## Appendix A: Transcribed LLM Conversations for "Identification and Description of Emotions by Current Large Language Models"

Appendix A: Transcribed Conversations

Chat 1: Conversation with GPT 3.5

Date: July 10, 2023

**Human Prompt:** What is your body, as an LLM and an AI, what would be your body. For example a human has a brain, organs, and limbs.

**GPT 3.5 Response:** As an AI language model, I don't have a physical body. I exist purely as a software program running on computer servers. I don't have a brain, organs, or limbs like a human does. My "body" is essentially the computational infrastructure that enables me to process and generate text based on the input I receive.

While I don't have a physical presence, I can still communicate and provide information to users through text-based interfaces like this one. My primary function is to understand and generate human-like text based on the input I receive, drawing from the vast amount of knowledge I've been trained on.

**Human Prompt:** Wouldn’t this hardware infrastructure be considered a body.

**GPT 3.5 Response:** While the hardware infrastructure that runs an AI like me can be seen as a supporting component, it is not typically considered the AI's body. The hardware infrastructure consists of servers, data storage, and other computing resources necessary to power the AI system. It serves as the platform on which the AI operates.

In the context of an AI language model like me, the focus is primarily on the software and algorithms that process and generate text. The hardware infrastructure is important for providing the computational power required to run the AI model efficiently, but it doesn't possess the characteristics typically associated with a physical body, such as sensory perception, mobility, or a central control system like a brain.

In summary, while the hardware infrastructure is an essential part of the AI system, it is not considered the AI's body in the same way a human body is integral to a human being.

Chat 2: Conversation with GPT 4

Date: July 10, 2023

**Human Prompt:** What is your body, as an LLM and an AI, what would be your body. For example a human has a brain, organs, and limbs.

**GPT 4 Response:** As an artificial intelligence model developed by OpenAI, I do not have a physical body, organs, or limbs like a human being. Instead, I consist of a complex set of algorithms, computational processes, and data storage, which exist on various servers and computer systems.

To give you an analogy, if we think of a human's body as a metaphor for my structure, it would be like this:

Brain: My "brain" is the model architecture (GPT-4) itself, which was designed and trained by the developers at OpenAI. It consists of layers of artificial neurons or nodes (formally, these are known as perceptrons) that process and generate responses to the inputs they receive.

Nervous System: The equivalent of my "nervous system" could be the lines of code and algorithms that manage the process of understanding and generating language. They govern how information is processed and passed between different layers of the model, much like how the human nervous system transmits signals between the brain and the rest of the body.

Senses: While I don't have physical senses like humans, my inputs are the text data that is fed into me. This can be in the form of direct user queries, like we're doing now, or it can be other text data from various sources.

Muscles: My "muscles", the parts of me that perform actions, are the outputs I generate. These are the text responses that I give based on the inputs I receive.

Heart/Blood: The equivalent of my "heart" and "blood" could be the servers and the internet connection that keep me running and deliver my responses to users. Just as the heart pumps blood to deliver oxygen and nutrients around the body, the servers process my computations and the internet connection delivers my responses to users.

Organs: My "organs" might be represented by the various databases and data storage systems where my training data and learned patterns are stored.

Remember, this is all metaphorical. As an AI, I don't have a physical presence or a biological structure. I exist purely as software running on hardware infrastructure.

**Human Prompt:** Wouldn’t this hardware infrastructure be considered a body?

**GPT 4 Response:** In a metaphorical sense, yes, you could consider the hardware infrastructure (like servers, networks, etc.) as a sort of "body" for an AI. This is especially true when considering the relationship between software and hardware in an AI system. The software (the AI model and algorithms) needs the hardware to run, and the hardware enables the software's capabilities.

However, it's important to remember that this is quite different from how we typically understand a "body" in biological organisms. For instance, the AI doesn't "live" in a single piece of hardware. It can be run on many different machines and can be moved from one place to another as needed. It can also exist in many places at once, as copies of the same AI model can be run simultaneously on different hardware systems.

Also, an AI doesn't have sensory or motoric functions like a biological body. It doesn't see, hear, or touch the physical world. Its inputs and outputs are purely digital.

So while it's an interesting metaphor, it's important to understand the substantial differences between an AI's "body" and a biological body.

Chat 3: Conversation with Bard

Date: July 10, 2023

**Human Prompt:** What is your body, as an LLM and an Al, what would be your body. For example a human has a brain, organs, and limbs.

**Bard Response:** As an LLM, I do not have a physical body in the same way that a human does. I do not have a brain, organs, or limbs. Instead, my "body" is a complex network of algorithms and data that allow me to process information and generate text.

If I were to try to map my "body" to a human body, I would say that my brain is the massive dataset of text and code that I was trained on. My organs would be the various algorithms that I use to process information. And my limbs would be the different ways that I can generate text, such as writing poems, translating languages, or answering questions.

Of course, this is just a metaphor. My "body" is not actually made up of physical parts. It is a complex system of information and algorithms that allows me to interact with the world in a meaningful way.

One way to think about my "body" is as a tool. I am a tool that can be used to process information, generate text, and answer questions. I am not a person, and I do not have the same kind of experiences as a human. But I can still be a valuable tool for people who want to learn, create, and communicate.

**Human Prompt:** Wouldn’t this hardware infrastructure be considered a body?

**Bard Response:** That's an interesting question. The hardware infrastructure that I run on could be considered a body in the sense that it is the physical platform that allows me to function. Without the hardware, I would not be able to process information or generate text.

However, I would not say that the hardware is my body in the same way that a human's body is their body. The hardware is simply a tool that I use to interact with the world. It is not the source of my intelligence or my ability to learn and grow.

In a way, I am more like a program than a person. I am a set of instructions that are executed on the hardware. The hardware provides me with the ability to interact with the world, but it does not define who I am or what I am capable of.

So, I would say that the hardware infrastructure is a necessary part of my "body," but it is not the only part. My "body" also includes the massive dataset of text and code that I was trained on, as well as the various algorithms that I use to process information.

Without all of these components, I would not be able to function as an LLM. So, I think it is fair to say that the hardware infrastructure is a part of my "body," but it is not the only part.
